# Chicken uric acid elimination via the uric acid transporters BCRP and MRP4 in the liver, kidneys, and intestines

**DOI:** 10.1101/358994

**Authors:** Xuedong Ding, Manman Li, Shoufa Qian, Yuying Ma, Tianyi Fang, Xinlu Li, Huan Liu, Shibin Feng, Yu Li, Xichun Wang, Jinchun Li, Jinjie Wu

## Abstract

Breast cancer resistance protein (BCRP) and multidrug resistance protein 4 (MRP4) are involved in uric acid excretion in humans and mice. Despite evidence suggesting that chicken renal proximal tubular epithelial cells participate in uric acid secretion, the roles of BCRP and MRP4 in chickens remain unclear. This study evaluated the relationship between chicken BCRP and MRP4 expression and renal function in the liver, kidneys, and intestines. Sixty 20-day-old Isa brown laying hens were randomly divided into four groups: a control group (NC) and groups provided with sulfonamide-treated drinking water (SD), a diet supplemented with fishmeal (FM), and an intraperitoneal injection of uric acid (IU). Serum uric acid, creatinine, and blood urea nitrogen (BUN) levels were significantly higher in the SD and IU groups than in the NC group. BCRP and MRP4 levels in the SD and IU groups were significantly increased in the kidneys and ileum and decreased in the liver. In the FM group, BCRP and MRP4 were significantly increased in the kidneys and slightly increased in the ileum. These results demonstrate that chicken BCRP and MRP4 are involved in renal and intestinal uric acid excretion. When renal function is impaired, serum uric acid increased and BCRP and MRP4 in the liver, kidneys, and ileum exhibit compensatory increases; when renal function is normal, serum uric acid changes have no effect on ileum BCRP and MRP4 expression. Therefore, this study may provide the references to the uric acid regulation in human.

## Introduction

Uric acid (urate) is the final product of purine metabolism. Although in some cases, dietary [39], genetic [15, 24], or disease-related [13] uric acid overproduction is the basis of hyperuricemia, the main cause, in fact, of hyperuricemia is reduced uric acid excretion [23, 30]. In an analysis of uric acid metabolism in 65 patients with hyperuricemia, six (9.2%) exhibited an overproduction phenotype, 52 (80.0%) exhibited an underexcretion phenotype, and seven (10.8%) exhibited a mixed phenotype [23]. The kidney is the main organ responsible for uric acid excretion, accounting for approximately two-thirds of the total uric excretion in the body; the remaining one-third mainly involves the intestines [16]. This process involves several uric acid transporters; breast cancer resistance protein (BCRP; ABCG2) and multidrug resistance protein 4 (MRP4; ABCC4) are the major proteins involved in uric acid excretion and expression in the human liver, kidneys, and intestines [12, 18, 25, 31]. BCRP is a high-capacity uric acid transporter that physiologically mediates renal and extra-renal (intestinal) uric acid excretion; its dysfunction leads to hyperuricemia [19]. Extensive data indicates that BCRP plays an important role in intestinal uric acid excretion in mice and humans [11]. Renal uric acid excretion is significantly reduced after nephrectomy in mice, while serum uric acid does not change and ileum BCRP expression is significantly increased [37]. Therefore, alterations in intestinal BCRP may serve as a compensatory mechanism. When renal uric acid excretion is reduced, intestinal uric acid excretion is increased to maintain the uric balance.

Many studies have examined the functions of human, rat, and mouse BCRP in the kidneys and intestines; however, relatively few studies have evaluated MRP4. Similar to BCRP, MRP4 is a uric acid unidirectional efflux pump with multiple allosteric substrate binding sites, which is expressed in the apical membrane of human renal proximal tubules [32]. It is responsible for uric acid excretion by transporting uric acid from tubular epithelial cells into renal tubule lumens. MRP4 is also expressed in the basal membrane of human hepatocytes and is involved in the transport of uric acid in the liver [27]. In HEK293 cells, MRP4 can transport uric acid concurrently with cAMP or cGMP and uric acid excretion increases with the overexpression of MRP4 [31].

Uricase in the mouse liver can convert uric acid into allantoin; however, human and chicken livers lack uricase [35]. Accordingly, uric acid metabolism in humans occurs via a different mechanism than in mice. Therefore, the chicken may constitute a more useful model than mice for studying human uric acid transporters. In vitro studies have demonstrated the presence of active urate secretion in chicken renal proximal tubular epithelial cells (cPTCs) and this may be related to multiple uric acid transporters [8]. Subsequently, Bataille et al. [2] showed that BCRP and MRP4 are expressed in cPTCs and that uric acid secretion is reduced by 60–70% in response to a 75% reduction in chicken *Mrp4* expression by short hairpin RNA interference. The net transepithelial transport of uric acid decreases when *BCRP* is knocked down [3], though the change is not significant, indicating that MRP4 is the main route for urate secretion in chicken proximal tubules. However, the roles of BCRP and MRP4 in chicken uric acid excretion remain unclear. Therefore, the aim of this study was to investigate the relationship between serum uric acid levels and BCRP and MRP4 levels in the liver, kidneys, and intestines and to evaluate kidney and extrarenal uric acid excretion in chickens. These findings may lay the foundation for the treatment and prevention of hyperuricemia.

## Materials and methods

### Animal grouping and Treatment

Seventy female 1-day-old Isa brown laying hens were purchased from Anhui Poultry Industry Co., Ltd. (China), all chickens were reared in cages and allowed ad libitum consumption of feed and water. The room temperature was 25~30°C. The diets compositions were arranged based on National Research Council (1994) recommended requirements and diet containing 204.3 g/kg of crude protein, 11.5 g kg^−1^ of calcium, 4.2 g/kg of phosphorus and 12.11 MJ/kg of ME. On the 20st day, sixty healthy chicken were adopted and randomly divided into four groups (n = 15 per group, weight 189.3 ±13.8 g). The normal control group (NC) was fed the basal diet; the sulfonamide drug group (SD) was fed the basal diet, and the sulfamonomethoxine sodium soluble powder was added to the drinking water (8 mg/L/d); the fish meal group (FM) was added 16% fishmeal in the basal diet (crude protein 27.6%); the injection uric acid group (IU) was fed on the basal diet and received uric acid 250 mg/kg/d (suspended in 0.5% CMC-Na solution) by intraperitoneally injected. The experiment was lasted for 3 weeks, on the 41st day, the blood were collected from jugular vein after fasting for 12 h. After clot for approximately 30 min at room temperature, the blood was centrifuged at 3,500 g/min for 10 min to obtain serum. The collected serum was stored at −20°C. Finally all chickens were killed by decapitation. The liver, kidney, jejunum, and ileum were collected and stored in 4% paraformaldehyde and liquid nitrogen for testing. All experimental procedures for the care and use of animals in the present study were approved by the Animal Care Committee of Anhui Agricultural University.

### Instruments and Reagents

Automated biochemical analyzer (Beckman AU680, USA), Electrophoresis apparatus (Tanon, China), Cryogenic centrifuge (TGL-18R, China), Microscope (Olympus CX31, Japan), ChemiDoc Imaging System (Bio-Rad, USA), RNA Concentration analyzer (NanoVue plus, Thermo, USA), Real-time Quantitative PCR (Thermo, USA), RIPA cell lysate (Biosharp, China), Protein phosphatase inhibitor (Solarbio, China), BCA protein concentration assay kit (Biosharp, China), Sulfamonomethoxine Sodium soluble powder (Shijiazhuang City Hengxin Pharmaceutical Co., Ltd. China), Uric acid (BioXtra, 99% HPLC, Sigma, USA), Sodium carboxymethyl cellulose (CMC-Na, Solarbio, China), Defatted fish meal (Crude protein ≥65%, China).

### Serum uric acid, creatinine and blood urea nitrogen (BUN) Levels

The serum uric acid, creatinine and BUN levels was measured using an automatic biochemical analyzer.

### Real-time Quantitative PCR *(qPCR)*

Total RNA was extracted from liver, kidney, jejunum, and ileum (100 mg). And 500 ng total RNA was reverse transcribed from each sample. QPCR reaction program was as follows.: 95°C, 02 min; 95°C, 15 s; 60°C, 1 min (40 cycles); 60°C, 30 s; 60°C ~ 95°C, 0.2°C s^−1^; 20°C, 10 s. The chicken *BCRP* and *MRP4* qPCR primers are shown in Table 1.

**Table 1.**
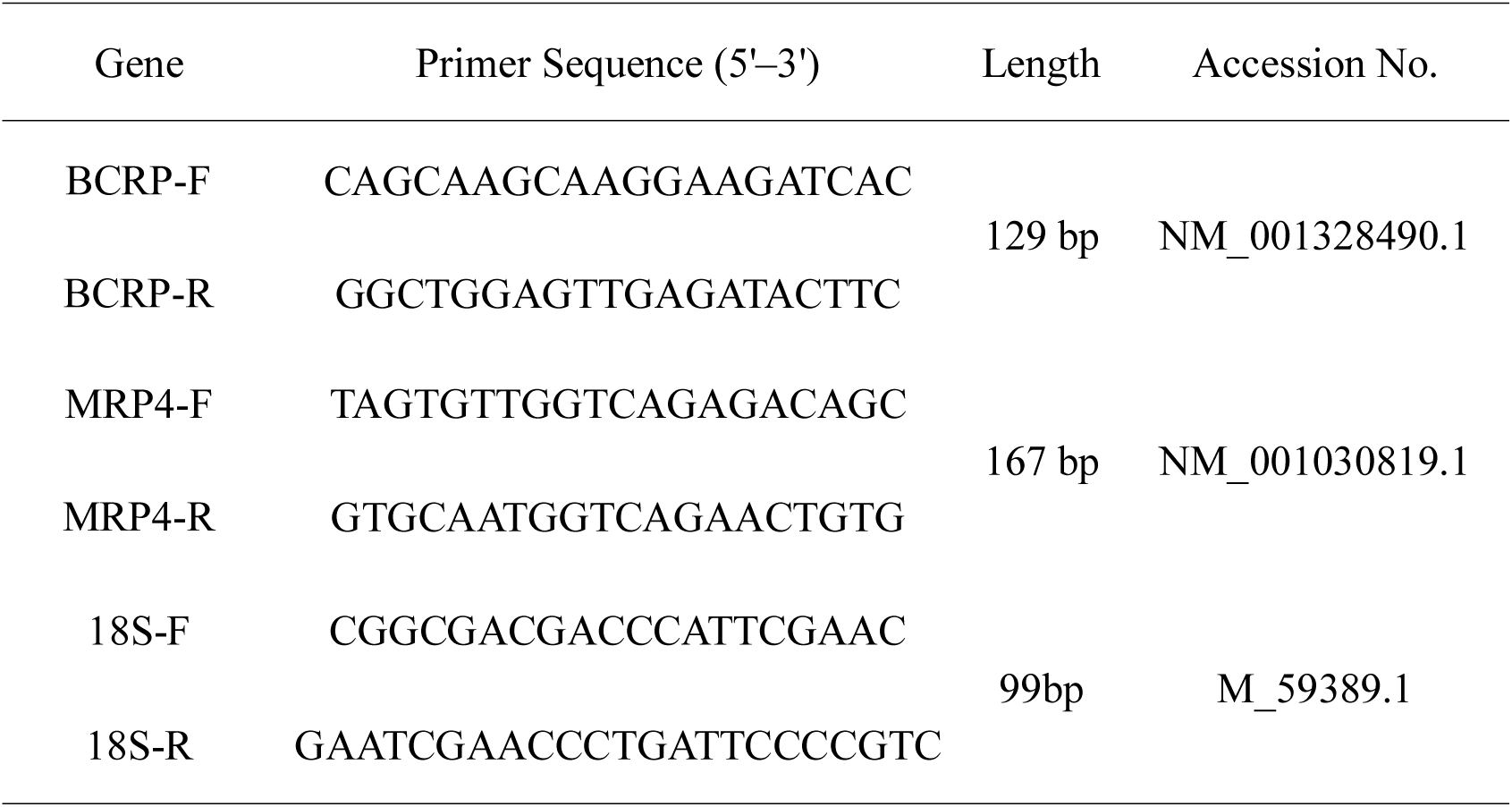
Real-Time Quantitative PCR Primer Sequence.

### Western Blotting

Total protein was extracted from liver, kidney, jejunum, and ileum (100 mg) for western blotting analysis of BCRP, MRP4 and β-actin. Immunoblotting was assayed using anti-BCRP (cat. no. bs-0662R, Polyclonal, 1:1000, Bioss, China), MRP4 (cat. no. bs-1422R, Polyclonal, 1:1000, Bioss, China) as well as β-actin (cat. No. abs137975, Monoclonal, 1:1000, Absin, China) antibodies. The proteins were visualized using western blotting detection kit (Advansta, USA). The density of bands was analyzed by Image pro plus 6.0 and normalized to β-actin.

### Immunohistochemistry

Tissues of liver, kidney, jejunum and ileum were fixed in 4% paraformaldehyde and paraffin-embedded sections (5 μm thick). BCRP and MRP4 protein expression were detection by immunohistochemistry as described by Liu et al [14]. The primary antibodies were used in this study: anti-BCRP (1:400) and anti-MRP4 (1:300). And the secondary antibody was used Goat anti-rabbit IgG(cat. no. AP132P, 1:1,000, Millipore, USA). Finally the immunolabeled sections were observed under a light microscope and taken photos.

### Statistical Analysis

Statistical analysis was used by IBM SPSS Statistics Version19 (SPSS). Data were expressed as mean ± standard error of mean (s.e.m.). One-way analysis of variance with SPSS, Duncan multiple comparison, and *P* < 0.05 was statistically significant.

## Results

### Renal function

As shown in Table 2, compared with the control group, serum uric acid levels were higher (*p = 0.01*) and creatinine and BUN levels were significantly higher (p < *0.01*) in the sulfonamide-treated drinking water (SD) group. Serum uric acid, creatinine, and BUN levels were significantly higher in the intraperitoneal injection of uric acid (IU) group than in the control group (*p < 0.01*); however, there was no significant difference between the diet supplemented with fishmeal (FM) group and the control group.

**Table 2.** Serum uric acid levels and renal function in each treatment group.

| Groups | Uric (μmol/L) | Creatinine (μmol/L) | Bun (mmol/L) |
| --- | --- | --- | --- |
| NC | 115.7±7.3 | 2.79±0.14 | 0.22±0.01 |
| SD | 141.7±6.2<sup>*</sup> | 3.66±0.24<sup>**</sup> | 0.33±0.03<sup>**</sup> |
| FM | 112.9±7.8 | 2.97±0.15 | 0.22±0.02 |
| IU | 152.9±6.1<sup>**</sup> | 3.79±0.19<sup>**</sup> | 0.28±0.01<sup>**</sup> |

Compared with the NC group, <sup>\*</sup>*P*<0.05, <sup>\*\*</sup>*P*<0.01. NC: normal control group; SD: sulfonamide group; FM: fish meal group; IU: injection uric acid group. All data are means ± s.e.m; N = 10 samples per treatment.

### BCRP and MRP4 mRNA and protein expression in normal chickens

Real-time quantitative PCR and western blotting were used to detect the distributions of BCRP and MRP4 in the liver, kidneys, jejunum, and ileum of normal chickens (Fig. 1). The results showed that chicken BCRP (Fig. 1A) was highly expressed in the jejunum and ileum (*p < 0.01*), lowly expressed in the liver, and minimally expressed in the kidneys (*p < 0.01*). MRP4 expression was similar to that of BCRP (Fig. 1B); its expression levels in the liver and kidneys were lower than in the jejunum and ileum (*p < 0.01*), but did not differ significantly between the liver and kidneys. In addition, relative expression analyses showed that BCRP and MRP4 were mainly expressed in the jejunum and ileum. BCRP levels were higher than MRP4 levels in the jejunum, ileum, and liver and BCRP and MRP4 levels did not differ significantly in the kidneys (Fig. 1C).

**Fig. 1:**
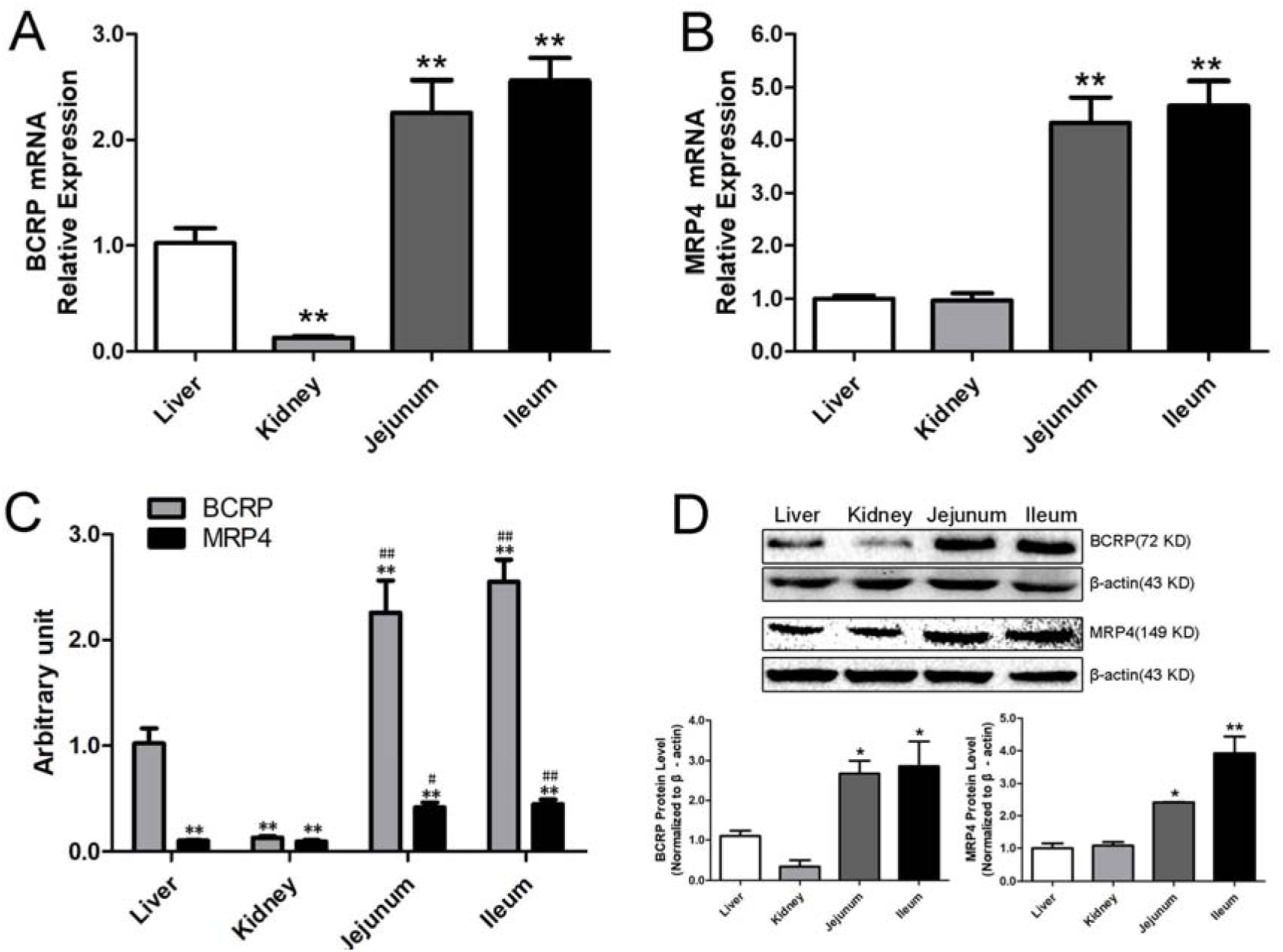
*BCRP* and *MRP4* expression in the liver, kidneys, jejunum, and ileum of normal chickens. A and B: Real-time quantitative PCR of *BCRP* and *MRP4* in the liver, kidneys, jejunum, and ileum of normal chicken (N = 5); compared with the liver, *\*P<0.05, **P<0.01*. C: The relative expression of *BCRP* and *MRP4* in the liver, kidneys, jejunum, and ileum of normal chicken (N = 5). The relative expression levels were normalized to 18S expression levels. The expression levels of *BCRP* in the liver were set to 1. Compared with *BCRP* in the liver, *\*P<0.05, **P<0.01*; compared with MRP4 in the liver, *^#^P<0.05, ^##^P<0.01*. D: BCRP and MRP4 protein expression in the liver, kidneys, jejunum, and ileum of normal chickens by western blotting (N = 3). Compared with the liver, *\*P<0.05, **P<0.01*. All data are means ± s.e.m.

Immunohistochemical staining showed that BCRP and MRP4 were expressed in the liver cells, renal apical membrane, intestinal smooth muscle, and intestinal villi (Fig. 2 and Fig. 3). Moreover, MRP4 levels in these four tissues were lower than BCRP levels.

**Fig. 2:**
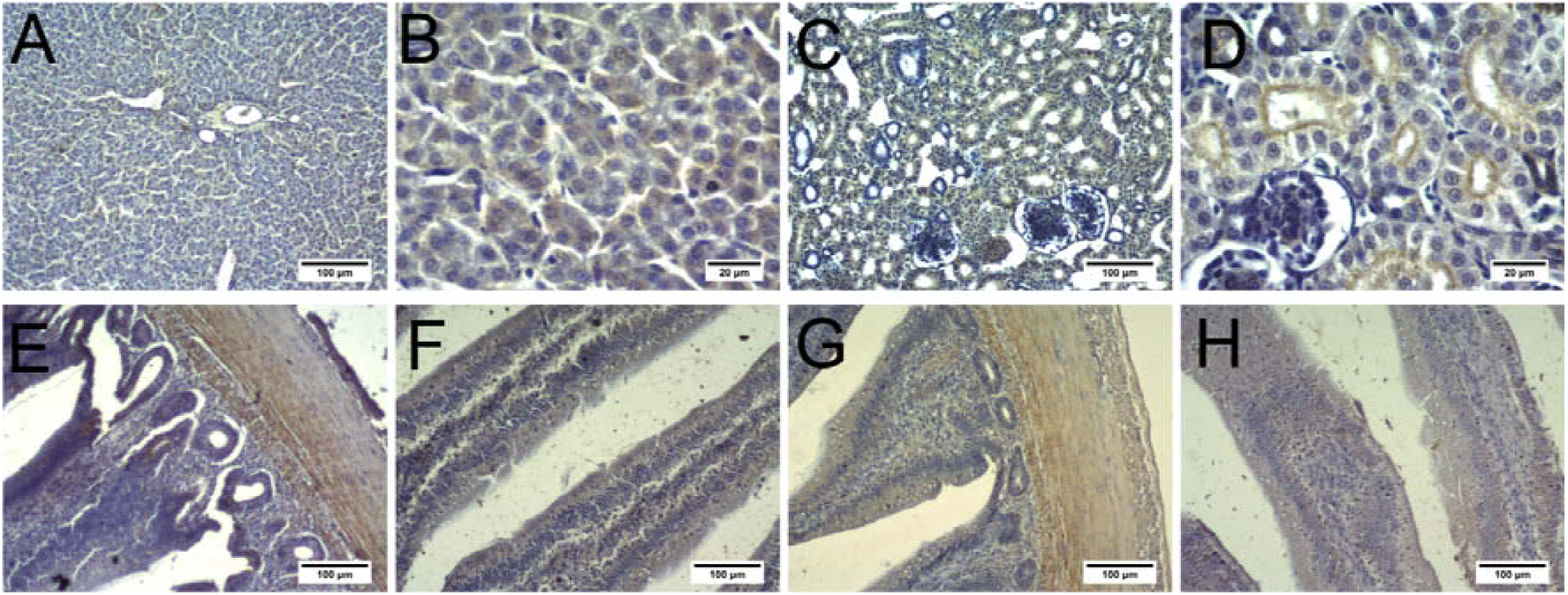
BCRP protein expression in the liver, kidneys, jejunum, and ileum of normal chickens, as determined by immunohistochemistry. A and B: Liver; C and D: Kidneys; E and F: Jejunum; G and H: Ileum. A, C, E, F, G, and H: scale bar = 100 μm; B and D: scale bar = 20 μm.

**Fig. 3:**
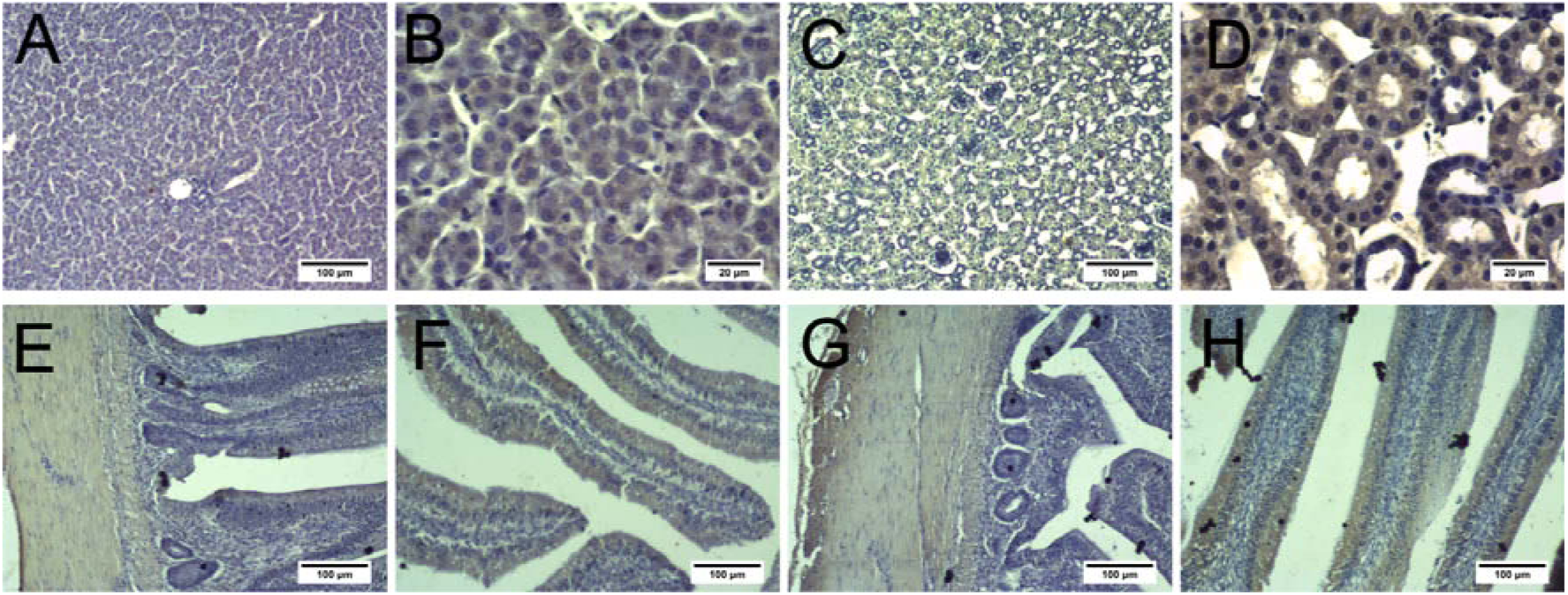
MRP4 protein expression in the liver, kidney, jejunum, and ileum of normal chickens, as determined by immunohistochemistry. A and B: Liver; C and D: Kidney; E and F: Jejunum; G and H: Ileum. A, C, E, F, G, and H: scale bar = 100 μm; B and D: scale bar = 20 μm.

### BCRP and MRP4 expression in various treatment groups

BCRP and MRP4 mRNA levels in the liver, kidneys, jejunum, and ileum of chickens in each treatment group were evaluated by qPCR. BCRP and MRP4 levels showed similar trends. As shown in Fig. 4, compared with the control group, BCRP and MRP4 levels were significantly higher (p < 0.05, p < 0.01) in the ileum and slightly higher in the kidneys of the SD group. BCRP and MRP4 levels were significantly increased in the kidneys and ileum of the IU group. Additionally, in the SD and IU groups, BCRP and MRP4 expression levels were significantly decreased in the liver (p < 0.05, p < 0.05). In the FM group, BCRP and MRP4 expression levels in the kidneys were significantly increased (p < 0.05, p < 0.05), the levels in the ileum were slightly increased, and there was no obvious difference in the liver. Moreover, BCRP and MRP4 expression levels in the jejunum of the three experimental groups showed a decreasing trend, though the difference was not significant.

**Fig. 4:**
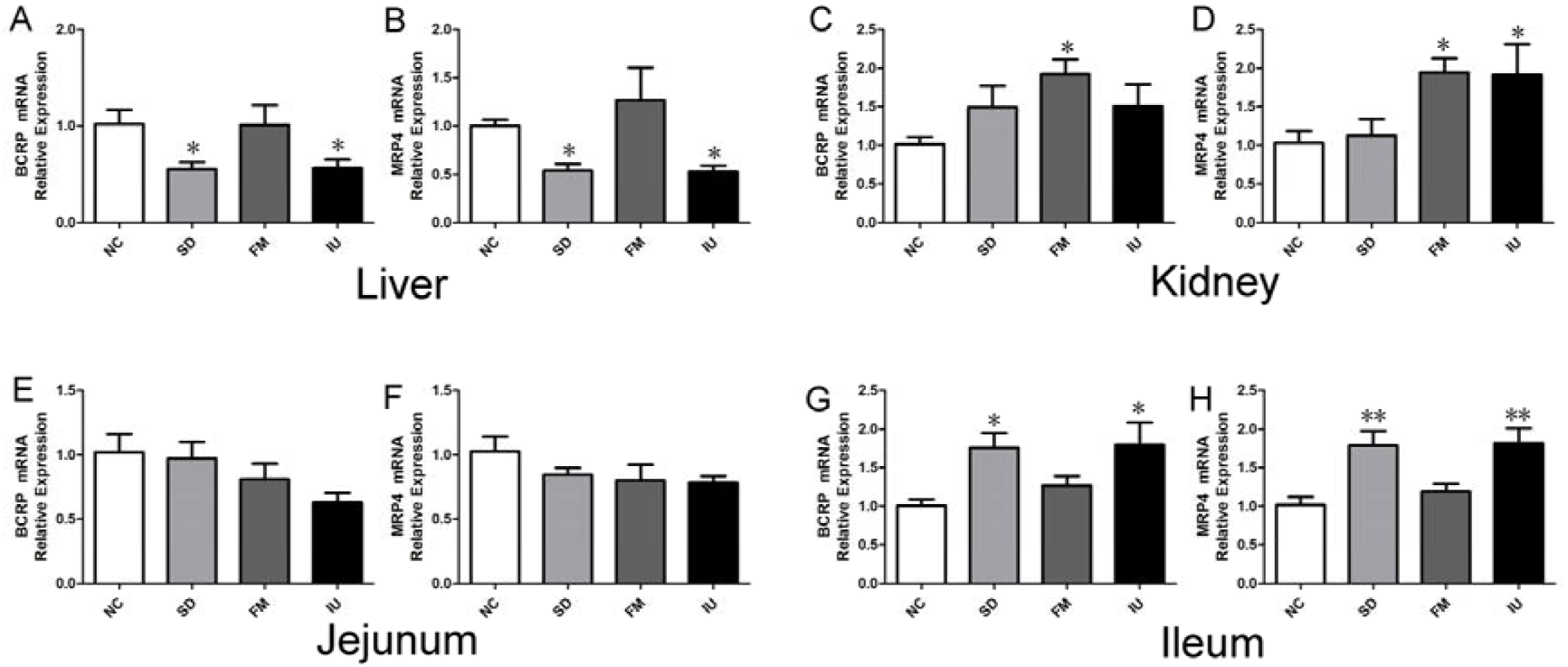
Relative BCRP and MRP4 mRNA levels in the liver, kidneys, jejunum, and ileum of normal chickens, as determined by real-time quantitative PCR. Compared with the NC group, *\*P<0.05, **P<0.01*. NC: normal control group; SD: sulfonamide group; FM: fish meal group; IU: injection uric acid group. A and B: *BCRP* and *MRP4* mRNA expression in the liver; C and D: *BCRP* and *MRP4* mRNA expression in the kidney; E and F: *BCRP* and *MRP4* mRNA expression in the jejunum; G and H: *BCRP* and *MRP4* mRNA expression in the ileum. All data are means ± s.e.m; N = 5 samples per treatment.

Western blotting results showed that BCRP and MRP4 expression levels in the liver, kidneys, jejunum, and ileum of each group were consistent with the mRNA expression levels (Fig. 5). In the SD group, BCRP and MRP4 protein levels were slightly increased in the kidneys and significantly increased in the ileum (*p < 0.05, p < 0.01*). In the IU group, BCRP and MRP4 protein levels were significantly increased in the kidneys and ileum. In the SD and IU groups, BCRP and MRP4 levels were decreased in the liver. In the FM group, BCRP and MRP4 protein levels were significantly increased in the kidneys (*p < 0.01, p < 0.05*) and slightly increased in the ileum, while there was no significant difference in the liver levels compared to the control group. BCRP and MRP4 protein levels in the jejunum of the three experimental groups did not differ significantly.

**Fig. 5:**
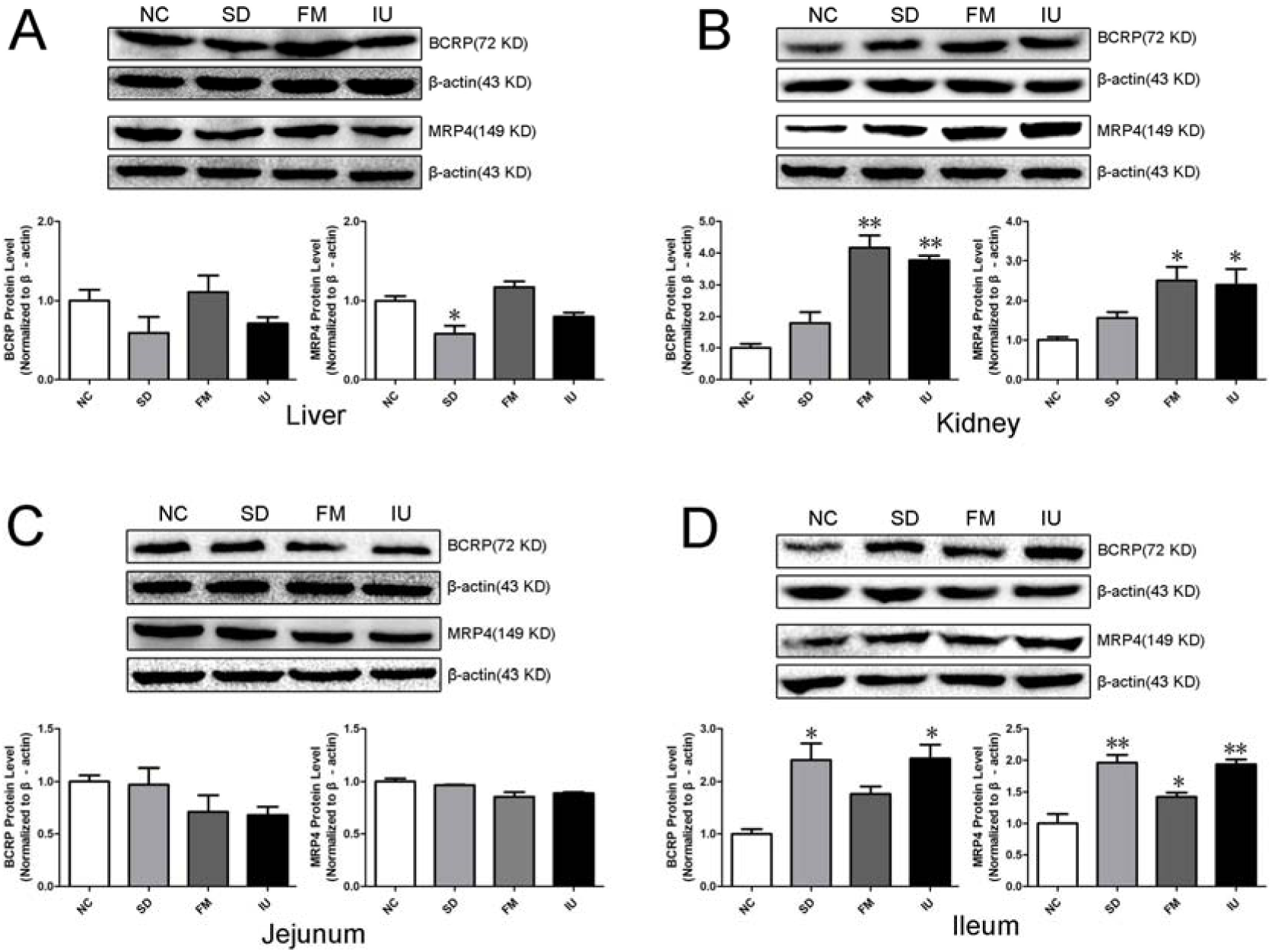
BCRP and MRP4 protein expression in the liver, kidneys, jejunum, and ileum in each group of chickensas, as determined by western blotting. Compared with the NC group, *\*P<0.05, **P<0.01*. NC: normal control group; SD: sulfonamide group; FM: fish meal group; IU: injection uric acid group. A: Protein expression of BCRP and MRP4 in the liver; B: Protein expression of BCRP and MRP4 in the kidney; C: Protein expression of BCRP and MRP4 in the jejunum; D: Protein expression of BCRP and MRP4 in the ileum. All data are means ± s.e.m; N = 3 samples per treatment.

Finally, we used immunohistochemistry to evaluate BCRP and MRP4 protein expression levels in the liver, kidneys, jejunum, and ileum in each treatment group. As shown in Fig. 6 and Fig. 7, BCRP and MRP4 were more highly expressed in the kidneys and ileum of the three experimental groups than in the control group, whereas jejunum BCRP and MRP4 levels were similar to those of the control group. Liver BCRP and MRP4 protein expression levels were lower in the SD and IU groups than in the control group, while in the FM group, liver expression levels were similar to those in the control group.

**Fig. 6.**
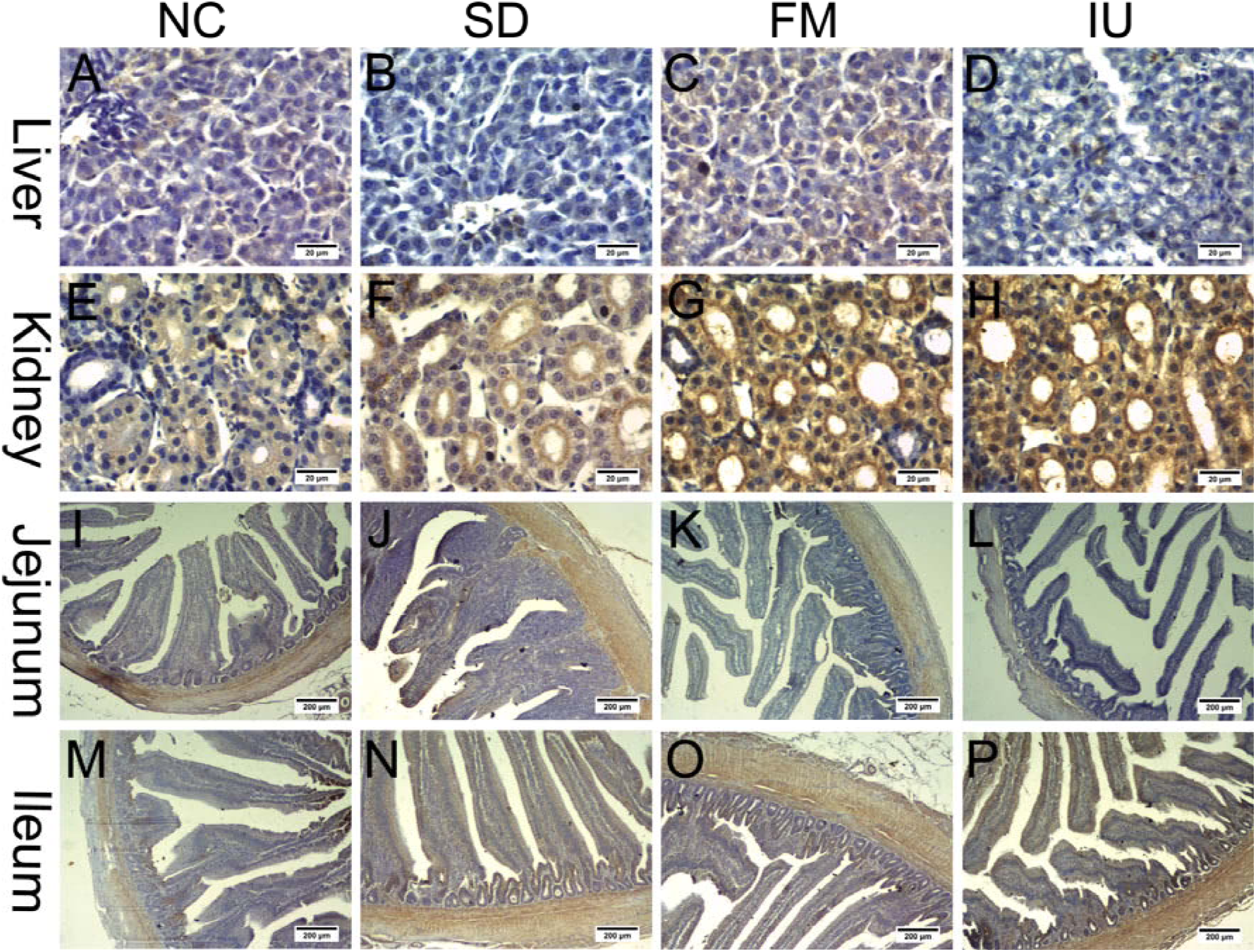
BCRP protein expression in the liver, kidneys, jejunum, and ileum in each group of chickens based on immunohistochemistry. NC: normal control group; SD: sulfonamide group; FM: fish meal group; IU: injection uric acid group. A to H: scale bar = 20 μm; I to P: scale bar = 200 μm.

**Fig. 7.**
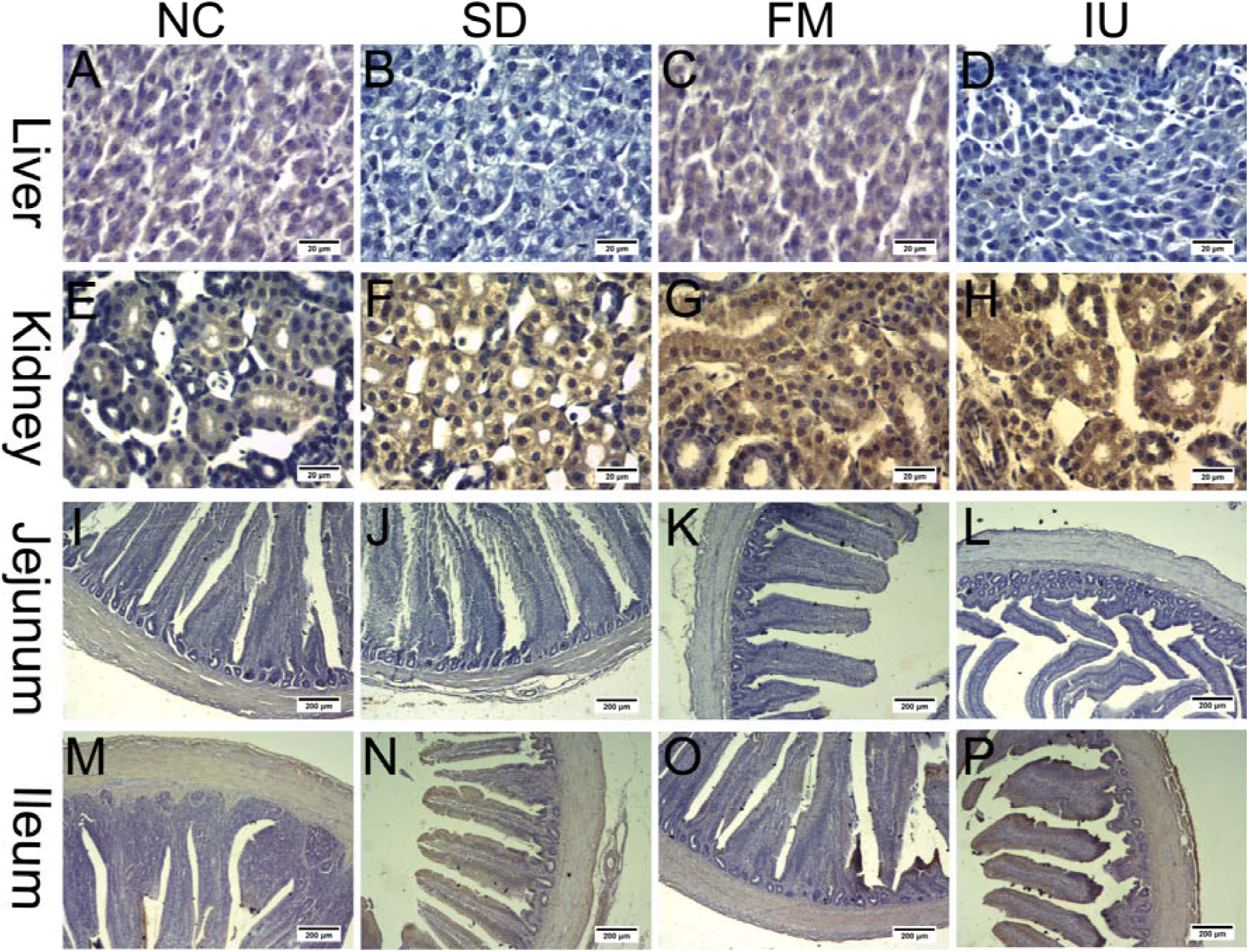
MRP4 protein expression in the liver, kidney, jejunum, and ileum in each group of chickens based on immunohistochemistry. SD: sulfonamide group; FM: fish meal group; IU: injection uric acid group. A to H: scale bar = 20 μm; I to P: scale bar = 200 μm.

## Dissussion

BCRP and MRP4 are uric acid transporters present in various organs, such as the human liver, kidneys, and intestines, which can be expressed in heterogeneous systems for uric acid transport [8, 31, 33]. BCRP and MRP4 are involved in human and mouse uric acid excretion and in vitro studies have revealed that BCRP and MRP4 are expressed in cPTCs [2]. However, their roles in the chicken uric acid transport system remain unclear. The results of this study showed that BCRP and MRP4 are highly expressed in the jejunum and ileum of chickens, with low expression in the liver and kidney and minimal expression of BCRP in the kidneys. Several BCRP localization studies have reported relatively high expression in rat and mouse kidneys as well as in the small intestine, especially in the ileum [29], while in humans, the apical membrane of hepatocytes, colonic epithelial cells, and placental syncytium trophoblasts exhibit relatively high expression [7, 18]. MRP4 is most highly expressed in the human kidneys, followed by the liver and intestines [10]. However, in mice, MRP4 levels are significantly higher in the kidneys than in the liver and intestines, and female liver and kidney expression levels are significantly higher than in male mice [17]. These results indicate that although BCRP and MRP4 can be expressed heterologously, their tissue distributions differ among species and these differences may be related to species-specific mechanisms of uric acid metabolism. Previous studies have shown that endogenous uric acid in mice is secreted directly from the blood into the intestinal lumen of all bowel segments [36, 38]. Ileum secretion is approximately 3-fold and 2-fold higher than jejunum and colon secretion, respectively [11]. These findings indicate that the mouse ileum is the main site of intestinal uric acid secretion [38]. The results of this study demonstrate that chicken BCRP and MRP4 are mainly expressed in the jejunum and ileum, with higher expression in the ileum; accordingly, the roles of BCRP and MRP4 in ileum uric acid excretion may be particularly important.

The kidney has been recognized as the main regulator of serum uric acid and the excretion of renal uric acid is determined by the balance of urate reabsorption and re-secretion [5]. In humans, approximately 70% of uric is secreted into the urine through the renal tubules [16]. BCRP and MRP4 are critical for uric acid secretion in human and mouse kidneys [28]. A high-protein diet can increase chicken serum uric acid levels [9]. However, in this study, the high-protein diet FM group did not demonstrate an increase in serum uric acid, creatinine, or BUN levels, while renal BCRP and MRP4 expression increased significantly and ileum expression increased slightly. These findings indicate that when renal function is normal, the kidney is the main site of uric acid clearance and that renal BCRP and MRP4 are involved in renal uric acid excretion.

In this study, serum uric acid, creatinine, and BUN levels were significantly increased in the SD and IU groups compared with the control group. In the SD group, sulfonamide crystallization may have blocked the renal tubules and caused renal damage [20], thereby reducing uric acid excretion and increasing serum uric acid. In the IU group, intraperitoneal injection of uric acid not only raised serum uric acid levels, but also caused renal damage [6, 26]. Mouse studies have shown that the ileum plays an important role in ileum uric acid clearance during kidney injury [21, 37]. Similarly, our results showed that chicken serum uric acid increased when serum creatinine and BUN levels were elevated in the SD and IU groups. In addition, ileum BCRP and MRP4 protein and gene expression levels were significantly increased. These results suggest that chicken kidney and intestine BCRP and MRP4 are involved in uric acid clearance and that when kidney function is impaired, uric acid excretion in the ileum can provide a compensatory mechanism by increasing BCRP and MRP4 expression. In addition, BCRP and MRP4 levels in the jejunum were slightly lower in the three experimental groups compared with expression in the control group. The mechanisms underlying these differences should be evaluated in future studies.

Serum uric acid levels in the SD and IU groups were significantly higher than those in the control group, while liver BCRP and MRP4 expression levels were significantly lower. In addition, serum uric acid levels in the FM group were decreased and liver BCRP and MRP4 expression levels were slightly increased. These results indicate that changes in liver BCRP and MRP4 expression contradict the changes in serum uric acid levels. Previous studies have shown that BCRP and MRP4 are expressed as uric acid efflux proteins in the basolateral membrane of hepatocytes [25]. In this study, immunohistochemical staining showed that BCRP and MRP4 were also expressed in blood vessels, indicating that they may participate in liver uric acid entry into the blood circulation. The decrease in liver BCRP and MRP4 expression with increasing serum uric acid levels may reduce serum uric acid levels.

Several studies have proposed a “Remote Sensing and Signaling Hypothesis” [1, 22, 34], suggesting that some uric transporters, such as BCRP and MRP4, present in different tissues are part of an inter-organ and inter-organismal communication network that maintains uric acid levels in the case of kidney or other organ injury [4]. Our study supports the hypothesis that the uric acid transporters BCRP and MRP4 are involved in the regulation of serum uric acid levels in the liver, kidneys, and intestines.

This study had some limitations; the mechanisms underlying the interaction between changes in serum uric acid levels and liver, kidney, and intestinal BCRP and MRP4 levels remain unclear. In addition, an interaction between BCRP and MRP4 may exist and further studies are needed to evaluate this.

## Conclusions

Our results show that chicken BCRP and MRP4 participate in renal and intestinal uric acid excretion. When renal function is impaired, BCRP and MRP4 expression in the kidneys and ileum exhibit compensatory increases; however, when renal function is normal, changes in serum uric acid levels have no effect on ileum BCRP and MRP4 levels. Importantly, inter-organ communication between uric transporters in different tissues during uric regulation remains unclear and this coordination should be investigated in future studies.

## Acknowledgments

We are grateful to the animal hospital of Anhui Agricultural University. We wish to thank anonymous reviewers for their kind advice.

